# Parasite transmission in aquatic ecosystems under climate change: joint effects of temperature, host behavior and elimination of parasite larvae by predators

**DOI:** 10.1101/769281

**Authors:** M. Gopko, E. Mironova, A. Pasternak, V. Mikheev, J. Taskinen

## Abstract

1. A moderate raise in temperature was suggested to enhance the impact of parasites on aquatic ecosystems. Under higher temperatures, poikilothermic animals (e.g. fish), increase their activity, which can result in a more frequent encounter with parasites. However, temperature increase may also trigger processes counteracting an increased risk of parasitic infections. For instance, removal of free-living stages of parasites by filter-feeding organisms can increase with temperature and potentially mitigate disease risk in ecosystems under climate change.
2. In our study, we aimed to find out whether an increased infection transmission under higher temperatures can be, at least, partly compensated by the increased removal of parasitic larvae be aquatic predators. In addition, we planned to reveal the behavioral mechanism underlying the more successful transmission of the parasite at higher temperatures.
3. We experimentally studied how temperature, the behavior of fish host (rainbow trout) and the presence of filter-feeding mussels in the environment influence transmission success of trematode larvae (*Diplostomum pseudospathaceum* cercariae) to fish host.
4. We found that temperature raise increased, while presence of filter-feeding mussels in the environment decreased infection intensities in fish. However, the effect of mussel’s presence was constant within the tested range of water temperatures (15-23ºC), which suggests that it cannot compensate for the observed increased transmission of parasites under temperature raise. The difference in fish individual behavior (swimming activity) before the exposure to parasites was a substantial factor the affecting host’s vulnerability to infection. However, fish motor activity only weakly correlated with temperature, therefore, it is unlikely to be responsible for the increased infection success under warmer conditions. After exposure to parasites, fish strongly decreased their activity. This decrease was temperature-dependent and more pronounced in bolder (more active) fish, which leads to lower variability in activity of fish exposed to parasites compared with the safe environment. Post-exposure activity did not influence the infection intensity.
5. In general, we showed that the elimination of trematode larvae by filter-feeders is unlikely to deter the potential effects of global warming on host-parasite interactions in temperate freshwater ecosystems.

## Introduction

Recent studies suggest that the impact of parasites on aquatic ecosystems can be considerably affected by climate change (Studer et al., 2010; Lõhmus & and Björklund, 2015; Marcogliese, 2016; Cable et al., 2017). In general, though it differs from one host-parasite system to another, a moderate increase in water temperature can enhance transmission of the majority of parasitic species, e.g. by increasing the rate and by lengthening of the annual period of larval production, affect life cycles and the global distribution of parasites (Harvell et al., 2002; Utaaker and Robertson, 2015; Lõhmus & Björklund, 2015; Barber et al., 2016; Baker et al., 2018; Mouritsen et al., 2018)◻. In addition, global warming causes multiple shifts in biology (growth, behavior, abundance, diversity, etc.) of their hosts and predators influencing interactions of these organisms with parasites (Macnab & Barber, 2011; Lõhmus & Björklund, 2015; Brunner & Eizaguirre, 2016)◻. Along with the fish-host immunity suppression caused by the temperature raise (Dittmar et al., 2014)◻, changes in the behavior of receptive hosts is one of the potential mechanisms providing increased parasite transmission. For instance, increased activity or ventilation rate (Pritchard et al., 2001; Mikheev et al., 2014; Lõhmus & Björklund, 2015) can lead to increased exposure of fish to parasites.

However, temperature raise may also launch ecosystem processes, which compensate for the increased transmission success of parasites. Free-living stages of parasites comprise a substantial share of the biomass in aquatic ecosystems (Lafferty et al., 2008; Kuris et al., 2008)◻ and many aquatic animals consume parasitic larvae, thus, significantly reducing transmission of parasites (Thieltges et al., 2008; Johnson et al., 2010; Welsh et al., 2014; Gopko et al., 2017). Feeding rates of ectothermic organisms are strongly temperature-dependent like most metabolic processes (Schmidt-Nielsen, 1997). For instance, removal of free-living stages of parasites by filter-feeders is suggested to increase with temperature up to a threshold level determined by physiological characteristics of predators (Burge et al., 2016)◻. However experimental data about the effect of temperature on the elimination of parasites by aquatic predators are still scarce (Goedknegt et al., 2015)◻ and do not include observations of host behavior. To our knowledge, there is only one study which reported that the presence of predators (barnacles) at higher temperatures has a stronger effect on infection transmission than at lower ones (Goedknegt et al., 2015)◻.

Change in fish vulnerability to infection caused by temperature raise could be mediated by fish behavior. For instance, under higher temperatures fish can increase their motor or/and ventilation activity which potentially increases exposure rate, thus increasing parasite’s chances to penetrate host skin and gills (Mikheev et al., 2014)◻. In addition, individual behavioral variation can also influence host vulnerability to infection. For instance, it was suggested that more risky and exploratory individuals (i.e. individuals with higher motor activity) might be at a higher risk of infection compared with shyer ones (Hoverman & Searle, 2016; Buck et al., 2018)◻. Though correlation between animal behavior traits and parasitic load was suggested in many studies (Hoverman & Searle, 2016; Barber et al., 2017; Cable et al., 2017)◻, an influence of individual’s personality on vulnerability to infection has rarely been tested experimentally (see however Koprivnikar et al. (2012) and Araujo et al. (2016)◻).

A recent study showed that filter-feeding freshwater mussels *Anodonta anatina* can significantly reduce transmission of the fish trematode *Diplostomum pseudospathaceum* by eliminating its free-living stages, i.e. cercariae (Gopko et al., 2017)◻. This parasite is very common in limnetic systems of temperate and boreal zones, infect a plethora of fishes, and can hamper fish farming (Valtonen & Gibson, 1997; Karvonen et al., 2006).

In the present study, we investigated the effect of temperature and mussels (*A. anatina*) on the transmission of a common fish trematode (eye fluke, *D. pseudospathaceum*) with a focus on potential interactions between these two factors and fish behavior.

Our main hypotheses were: (1) fish (*Onchorhyncus mykiss*) will be more vulnerable to parasitic infection under higher temperature due to increased activity; (2) mussels can remove trematode cercariae from the water in a wide range of temperatures and their impact on the reduction of the infection in fish is temperature-dependent (i.e. they can at least partly compensate for increased vulnerability to parasites in fish caused by a temperature raise).

## Material and methods

### Study objects

All experiments were conducted at the Konnevesi research station (University of Jyväskylä) in summer 2017. We used a common fish trematode *D. pseudospathaceum* as the parasite, rainbow trout *O. mykiss* as the host and freshwater mussels *A. anatina* as predators of cercariae.

The eye fluke *D. pseudospathaceum* has three hosts in its life-cycle: freshwater mollusks (the first intermediate host), different fishes (the second intermediate host) and fish-eating birds as definitive hosts (Valtonen & Gibson, 1997; Karvonen et al., 2006)◻. In fish, this parasite localizes in the eye lenses and decreases host fitness by impairing vision (Owen et al., 1993; Karvonen et al., 2004a)◻ and manipulating host’s behavior (Seppälä et al., 2004; Mikheev et al., 2010; Gopko et al., 2015, 2017a)◻. Young-of-the-year rainbow trout were obtained from a commercial fish farm and acclimated in the laboratory at least for two weeks before the experiments. At the fish farm, rainbow trout were maintained in ground water and, therefore, were free of macroparasites. *A. anatina* mussels were collected from Lake Jyväsjärvi and were acclimated at the lab for a week before the experiments. Each mussel was observed to filter actively (siphons protruded) before the start of the experiment. Infected pond snails *Lymnaea stagnalis* collected from Lake Konnevesi were used as a source of *D. pseudospathaceum* cercariae. The shedding of cercariae by snails was checked visually by incubation of snails in glasses with filtered lake water under the bright light for several hours. Since in Finland (including Lake Konnevesi) *L. stagnalis* is typically infected with *D. pseudospathaceum* rather than other related diplostomidae species (Louhi et al., 2010; Rellstab et al., 2011), the cercariae were identified microscopically by their morphology.

### Experimental design

The experiment was divided on ‘tests’ conducted seven times in a row (at different temperatures) so that each test was started after the previous one ended. In each test, fish randomly chosen from the stock maintained in the laboratory were placed individually in 26-28 white containers (30×40×25 cm) filled with 12L of filtered lake water and were acclimated for an hour before exposure to cercariae. Fish were randomly assigned to four treatments (6-7 replicates in each). During the acclimation period, water in half of the containers was slowly warmed with aquarium heaters, while in another half, similar heaters were placed, but switched off. In addition, in half of the containers in each heating treatment, we placed live *Anodonta anatina* (one mussel per container), while in other half closed empty shells of mussels. Empty shells and switched off heaters were placed in containers to minimize the difference in fish behavior between the treatments (Gopko et al., 2017b)◻. Therefore, there were the following four treatments (6-7 fish in each): (1) containers with heating and the presence of live mussel (H+M+), (2) containers with heating and the presence of empty shell, i.e. ‘mussel’ control (H+M-), (3) containers with switched-off heaters and live mussels (H-M+) and (4) with switched-off heaters and empty shells (H-M-).

Tests were started at the same time of the day (between 0:30 and 1:30 p.m) to exclude potential effects of the circadian rhythms. Each test lasted for two days (the first day – infection, the second – dissection). In three tests, the temperature in containers with heating was set close to 19.5°C (mean±SD = 19.6±1.59°C), while in four others it was around 22.5°C (22.6±1.48°C). In control containers, the temperature was about 15-17°C (16.0±0.70°C). These values are typical of the surface layer in Lake Konnevesi after wind mixing in summer (mean daily temperature range 11.9–20.0, mean±SE = 16.1±1.23°C) (Kuha et al., 2016). Thus, the lowest water temperature in our experiment reflected natural conditions in nearshore regions of this lake. In addition, these temperatures are also similar to mean summer temperatures in temperate lakes (mean±SE = 16.8±0.52°C), which were calculated using data from ‘laketemps’ package (Sharma et al., 2015)◻ (see Supplement, Methods 1, for details). Therefore, temperatures in containers with heating reflect moderate predictions of temperature increase (1 – 5°C) by the end of the 21st century (IPCC, 2014, 2014) being far from the most pessimistic and extreme predictions for the temperate lakes in the northern hemisphere (Sharma et al., 2007)◻.

The temperature was measured in each container before the first fish activity tracking (see below) and at the end of the experiments (after removing fish from containers), and did not change significantly during this period. Temperature values obtained during post-experimental measurement were used in the statistical analysis.

However, there was a substantial temperature variation both among controls in different tests (due to changes in the outside temperature) and among heated containers, because our heaters cannot be precisely calibrated. Therefore, in the statistical analysis, we treated temperature as a continuous predictor, while statistical models, where the temperature was considered as a factor are presented in a Supplement (Methods 4, Results).

In total, 180 fish were used in the statistical analysis (see, however, *Fish activity tracking* section), because 16 individuals were lost due to jumping out from containers, death for unknown reasons or obvious signs of sickness and therefore were excluded from the sample. Fish loss never exceeded 3 individuals per test and the resulting number of rainbow trout used in all treatments were similar ranging from 43 to 47 fish. Therefore, it is unlikely that an uneven fish loss in different treatments can influence the results of the statistical analysis.

### Infection protocol and dissections

Fish were exposed to freshly produced *D. pseudospathaceum* cercariae obtained from five *L. stagnalis* snails less than 2 h before the exposure. The infection dose was 300 cercariae per fish, the exposure time was two hours.

After each test, rainbow trout were caught and placed individually in 8L flow-through tanks for 24 hours to let parasites reach eye lenses of the fish. Then fish were killed with an overdose of MS222, weighted and dissected. The number of *D. pseudospathaceum* metacercariae in the eye lenses of the fish was counted using a dissection microscope (32× magnification).

### Fish activity tracking

We video recorded fish behavior at different temperatures from above the aquaria for 5 minutes before and after exposure to parasites (one hour after the addition of cercariae). A grid (10 × 10 cm) was drawn on the bottom of each test tank and activity was measured as a number of gridlines crossed by fish in a 5-minute interval. Records were analyzed blindly (i.e. investigator was unaware about the treatment to which an observed fish belonged). Cameras were switched on from outside to avoid the influence of the investigator on fish behavior.

Unfortunately, due to a technical problem, all videos from one of the tests from the mild heating treatment were lost. In addition, several records were excluded from the sample, because some containers were partly out of camera range. Therefore, activity video records were obtained only for 142 fish.

### Statistical analysis

#### Influence of environmental conditions and fish weight on the infection intensity

Linear mixed models were used to estimate the influence of temperature and presence/absence of alive mussel in the environment. The practical and widely used strategy to find out which variables should be included in the model is a step-down (backward) model selection, however, its too straightforward implementation (i.e. including too large a set of possibilities) can turn into a data-dredging (Bolker, 2007, p. 277; Kuznetsova et al., 2017)◻. Therefore, we first formulated a biologically sensible model of interest, where all variables and the interaction purposefully tested in our study were included, and then simplified the model using backward selection tool from the ‘lmerTest’ package (Kuznetsova et al., 2017)◻.

The model was the following: log(infection intensity) ~ fish mass (covariate) + temperature (covariate) + alive mussel presence/absence (factor) + temperature*alive mussel presence/absence + experiment identity (random factor). Since we were interested in certain double interaction (temperature*alive mussel presence/absence) we included only this double interaction in our model of interest. The response variable (infection intensity, i.e. the number of *D. pseudospathaceum* metacercariae in fish) was log-transformed to meet model assumptions. To verify that we did not miss some important interactions, we also tested the model including all possible interactions using a similar approach. The resulted models were identical (see results), which suggests that models with higher order interactions are unlikely to explain the data substantially better than the model obtained by the model of interest simplification.

To account for the influence of fish activity on its vulnerability to parasites, we used an abridged dataset, since recordings of rainbow trout behavior were not available for all fishes (see the explanation in the *Fish activity tracking* section) and, therefore, in this case, fish activity was included in the model of interest. In all other respects, the statistical analysis was similar to the described above. We created two separate sets of models for fish activity before and after the exposure to cercariae. P-values were calculated using Kenward – Roger’s procedure for the approximation of degrees of freedom implemented in lmerTest package (Kuznetsova et al., 2017)◻.

To present the results of the mixed-effect models graphically, partial regression plots were drawn (see the details in the Supplement, Methods 2).

### Activity

A paired t-test was used to compare fish activity before and after the exposure to parasites. We used Fligner-Killeen test of homogeneity of variances, which is suggested to be robust against the departure from normality (Conover et al., 1981)◻ to compare, whether variation in fish activity before the exposure to parasites was larger compared with variation after the exposure. The robust test was chosen, since the data on post-exposure fish activity violated the normality assumption (Shapiro – Wilk’s test: W = 0.97, p-value = 0.005).

To check whether environmental conditions influence fish activity before and after addition of cercariae in the containers, we started with linear mixed models where fish activity before the exposure to parasites, fish activity after the exposure and differences in activities before and after exposure served the response variables and experiment ID was a random factor. Presence of the alive mussel in the container, temperature, fish mass and interactions between the variables were components of the full model, which was then simplified using a backward selection. However, an addition of the random factor did not appear to explain a substantial amount of variance in these models (p > 0.3 in both cases). Therefore, the random effect was deleted from the models and we proceeded with simple general linear models. The variable ‘temperature’ was centered by subtracting the mean to make the estimates of regression coefficients more biologically sensible.

We also checked whether fish with a different baseline level of activity (i.e. activity before adding cercriae) differed in their reaction to the presence of parasites in the environment (i.e. change in activity after adding cercariae). More technically speaking, we regressed fish pre-exposure activity versus the difference between pre- and post-exposure activity. However, the statistical evaluation of such a relationship is usually complicated because of two methodological concerns known as regression to the mean and mathematical coupling (Hayes, 1998; Tu & Gilthorpe, 2007)◻. To account for these problems, we used a method proposed by Tu et al. (2005)◻. For certain formulas and details see the Supplement (Methods 3), however, in brief, we calculated a correlation coefficient between pre-exposure activity and difference between pre- and post-exposure activity.

Then, a correct null hypothesis was determined taking into account the correlation between pre- and post-exposure activity of fish and mathematical coupling. Finally, both observed and expected (null hypothesis) correlation coefficients were *z*-transformed and a difference between them was compared with 0 using the *z*-test.

The models, where the temperature was considered a categorical variable (three heating treatments) were also fitted (see the Supplement, Results). Their results were very similar to the ones presented in the main text of the article.

All statistical tests were performed using R (R Core team, 2018)◻. A package ‘lme4’ (Bates et al., 2015)◻ was used to fit linear mixed models and get estimates of the regression coefficients. ‘ggplot2’ (Wickham, 2009)◻ and ‘sjPlot’ (Ludecke, 2018)◻ packages were utilized to visualize the data.

## Results

### Infection intensity

Mean (±*SE*) fish weight constituted 7.77±0.15 g (total 180 ind.) and 7.48±0.15 g (142 ind. of the abridged “activity dataset”). Fish size did not differ between the treatments (with alive mussel vs control) both for full and abridged datasets (ANOVA: *F_1, 178_* = 0.13*, p* = 0.72 and *F_1, 140_* = 0.12*, p* = 0.74 respectively). Mean±SE infection intensity was 46.8±2.41 in the full and 39.2±2.12 metacercariae per fish in the abridged dataset.

Linear mixed models comparison (i.e. procedure of backward model selection) showed that adding interaction terms did not lead to significant improvement of the model fit. Importantly, the interaction between temperature and presence of alive mussels was non-significant (*F_1, ‘173.1_* = 1.50, *p* = 0.22, see also Table 1a), which suggests that the ability of *A. anatina* to eliminate cercariae does not change substantially under the tested temperature. Moreover, adding one of the main effects to the model (fish mass) also did not significantly increase the amount of variance explained by the model (Table 1a). However, we decided to keep this predictor in the final model, since it seems biologically relevant and important. When mass was excluded from the model, p-values related to other predictor variables and the magnitude of estimated coefficients did not change substantially. Therefore, the final model contained only the main effects and test ID (random effect) as predictors (Table 1a). It showed that the effect of heating was significant and there was a 1.094-fold (exp(0.09) = 1.094) increase in parasitic load per each additional 1°C (Table 1a, Fig. 1A, C). The presence of the alive mussel in the environment decreased the *D. pseudospathaceum* infection intensity in fish by ~28% (Table 1a, Fig. 1A, C).

**Table 1.** GLMM on full and abridged datasets summary tables. Log-transformed infection intensity is a response variable.

|  | a. Full dataset models |  |  |  |  | b. Abridged dataset models |  |  |  |  |
| --- | --- | --- | --- | --- | --- | --- | --- | --- | --- | --- |
| Fixed effects | df | *t* | *p*-value | Est. | SE | df | *t* | *p*-value | Est. | SE |
| + temperature | 173.8 | 9.00 | <**0.0001** | 0.090 | 0.010 | 132.1 | 7.43 | <**0.0001** | 0.085 | 0.011 |
| + alive mussel | 173.0 | -5.46 | <**0.0001** | -0.329 | 0.060 | 132.0 | -4.71 | <**0.0001** | -0.327 | 0.069 |
| + activity (before) |  |  |  |  |  | 132.3 | 2.14 | **0.034** | 0.0014 | 0.0007 |
| + fish mass | 173.9 | 1.50 | 0.13 | 0.025 | 0.016 | 132.4 | 0.95 | 0.35 | 0.019 | 0.020 |

**Fig. 1.**
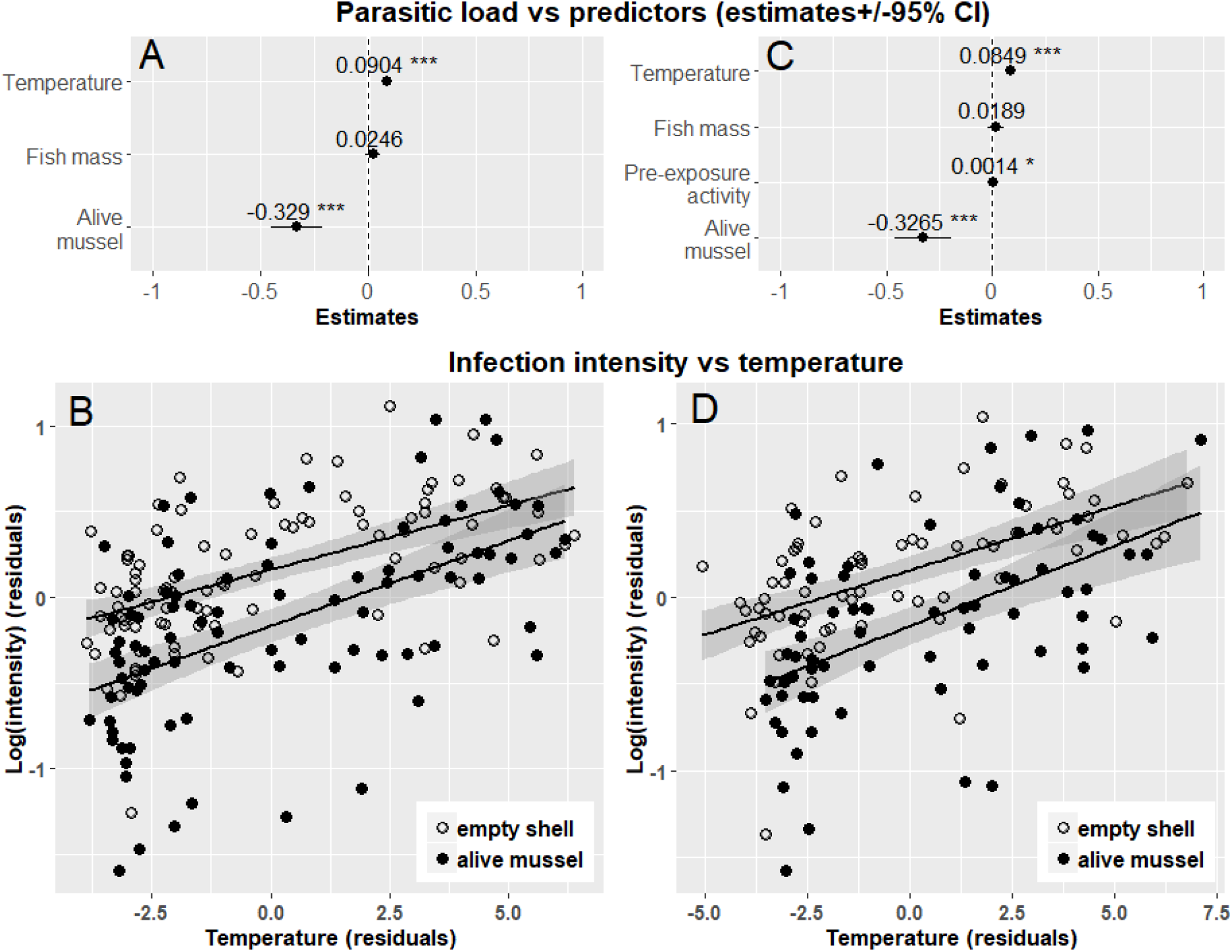
Regression coefficients plots (A, C) and partial regression plots (B, D) showing the influence of the temperature on the infection intensity in rainbow trout for the models fitted on the full (A, B) and abridged dataset (C, D). In both cases, the presence of alive mussel in the container caused a substantial decrease in the infection intensity in fish, while temperature increase led to higher infection intensities. The regression lines for containers with alive mussels and control containers are almost parallel, confirming the lack of interaction between the temperature and presence of alive mussel in the environment. Fish, which were more active prior to the exposure, were more infected compared with less active fish (A, C) (about 15% increase in the infection intensity per 100 additional lines crossed by fish in five min).

In the set of models, where the fish activity was included, the results were similar. Interactions were also not significant and were excluded from the final model. The effect of mass was again non-significant, however, this predictor was left in the model for its biological relevance, as described above. The effect of temperature was still highly significant (see table 1b, Fig. 1B, D). Fish activity before the exposure varied substantially among fish (with range 0-278 and mean±SE = 117.8±4.75 crossed lines/5 min). The effect of fish pre-exposure activity on the infection intensity was relatively week (about 1.0015 increase of infection intensity per each additional gridline crossed by fish). Interestingly, when fish activity after exposure (range 1-155 and mean±SE = 61.4±3.21 lines/5 min) was added to the model instead of pre-exposure activity, it had no significant effect on the infection intensity (*t*_132.2_ = 0.67, *p* = 0.51). The difference in fish activity before and after exposure to cercariae also was not a significant predictor of the infection intensity (*t*_132.5_ = −1.77, *p* = 0.14).

When the temperature was added in the model as a factorial variable, the results were very similar to the presented above (see the Supplement, Fig. S1 and Table S1).

### Activity

Paired t-test showed that before the exposure to parasites, fish were significantly more active than after the exposure (*t_141_* = 10.5, *p* < 0.0001, Fig. 2A, B). Moreover, fish were more variable in activity levels before the exposure to parasites (Fligner-Killeen test: χ^*2*^ = 12.43, *p* = 0.0004, Fig. 2A, B). Interestingly, there was only a weak correlation between fish activity before and after exposure (Spearman’s rho: r_s_ = 0.19, p = 0.03).

**Fig. 2.**
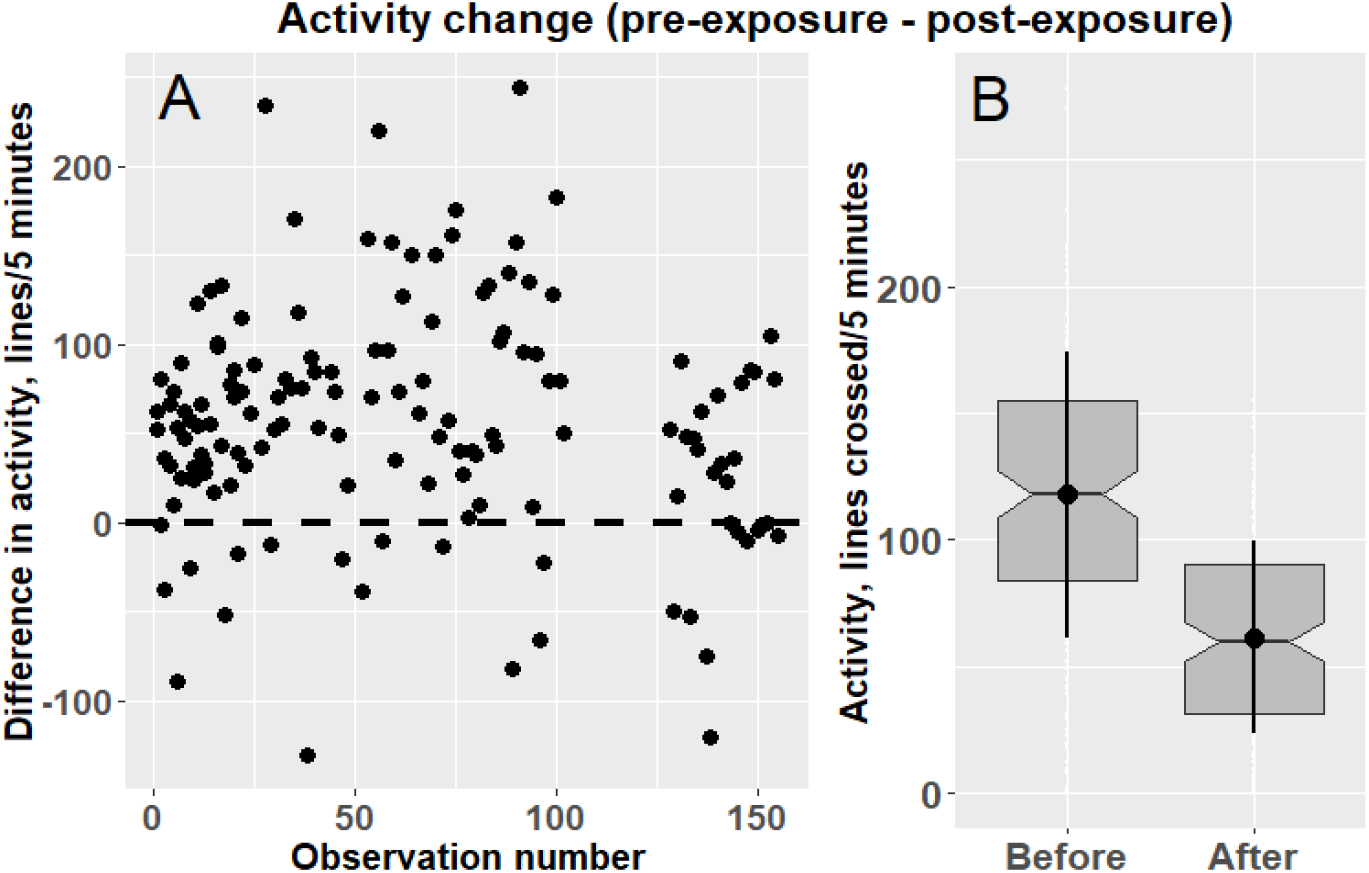
Fish activity before and after the exposure to parasites. A. Difference in activity (pre-exposure activity minus post-exposure activity). Before the exposure to cercariae, most of the fish were more active than after the exposure (dots above the dashed zero line vs triangles below it). B. Fish activity and between individual variance in activity before the exposure was significantly higher than after it. The box on the plot represents the median with the interquartile range (IQR). The notches represent roughly 95% confidence intervals (1.58*IQR/sqrt(N)) for the medians. Dots with whiskers are mean±SD fish activity.

For the pre-exposure activity, we found that the model, where only the temperature was a predictor, fits our data significantly better than only intercept model (*F_1, 141_* = 8.49, p = 0.004), while addition of other predictors and interactions did not explain the significant additional amount of variance. Fish activity increased with the temperature increase (Fig 3A) by extra four lines per each additional 1°C (Estimate±SE = 4.27±1.47). However, when the interaction between temperature and presence of alive mussel along with both main effects was included in the model its contribution was marginally significant (*F_1, 139_* = 3.63, p = 0.059, Fig. 3A). In other words, fish in both treatments became more active with increasing temperature. In the presence of alive mussels, they tended to increase activity even more.

**Fig. 3.**
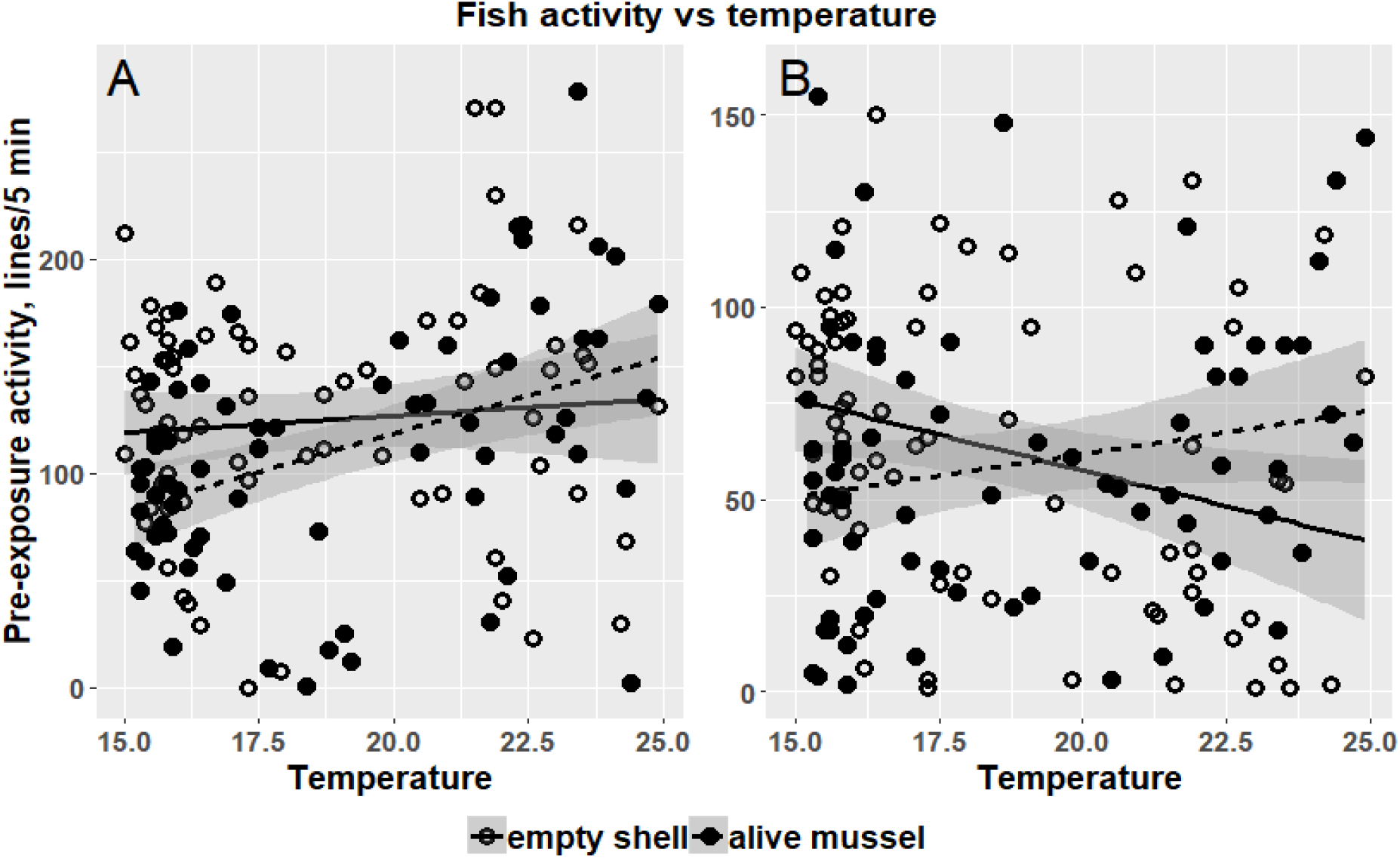
Rainbow trout pre-exposure (A) and post-exposure (B) activity depending on temperature. (A) Before the exposure to parasites, fish activity increased with temperature both in the presence of the alive mussel and in the control (empty mussel shell) (Estimate±SE = 4.27±1.47 for the similar slopes model). Though the slopes of regression lines for both groups did not differ significantly (*p* = 0.06), the line for fish in the presence of alive mussels was steeper (dashed line). (B) After exposure, the slopes of regression lines became significantly different. In the presence of alive mussels, fish still increased their activity with increasing temperature, while in the control, fish decreased their activity with temperature.

For the post-exposure activity, the model including the presence/absence of alive mussel in the container, temperature, and interactions of these effects was found the most parsimonious one. There was a significant effect of temperature (Estimate±SE = −3.71±1.45, *t* = −2.57, *p* = 0.011) and interaction between the temperature and presence of alive mussel in the model (Estimate±SE = 5.95±2.00, *t* = 2.97, *p* = 0.004 Fig. 3B). It means that in the containers with alive mussels fish post-exposure activity increased with temperature (regression coefficients was −3.71 + 5.95 = 2.24), while in containers with empty shells fish activity even decreased with temperature raise, and the slopes of the regression lines differ significantly between the treatments. Though we found a significant influence of temperature on fish pre- and post-exposure activity in our models, the amount of variance explained by our predictors was fairly small (6% and 7% respectively).

Temperature influenced the degree of activity change after the exposure to cercariae, i.e. pre-exposure activity minus post-exposure activity (Estimate±SE = 4.95±1.67, *t* = 2.97, p = 0.004, Fig. 4), while the addition of treatment (alive mussel/empty shell) and the interaction in the model did not increase additional amount of variance (Fig. 4). Fish changed their activity more under high temperatures compared with low temperatures.

**Fig. 4.**
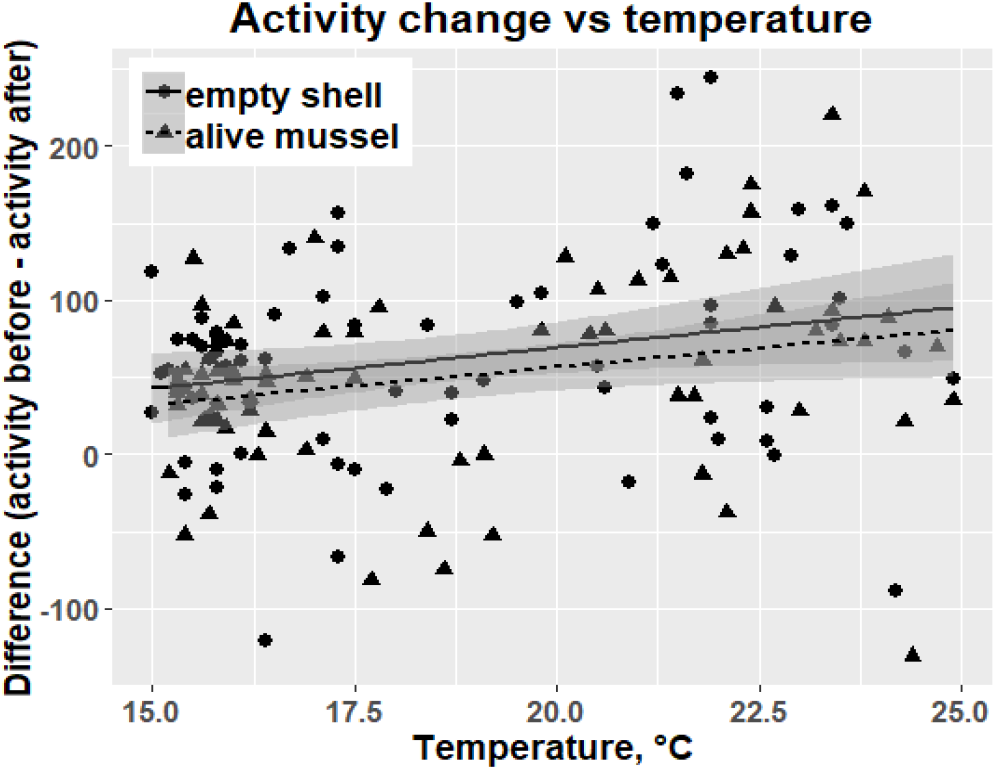
The decrease in fish activity after the exposure to parasites was temperature dependent and more prominent at higher than at lower temperatures (extra 4.4 lines crossed per each additional 1ºC). However, this effect was not modified by the presence/absence of alive mussels (regression lines are almost parallel with overlapping confidence intervals).

Fish more active before the exposure to parasites decreased their activity stronger compared with less active individuals (Fig. 5). The coefficient of correlation between baseline value (pre-exposure activity) and activity change was *r* = 0.81, which is significantly (*z* = 3.86, *p* = 0.0001) higher than null-hypothesis value (r = 0.66) calculated following Tu et al., 2005 (See Methods and Supplement).

**Fig. 5.**
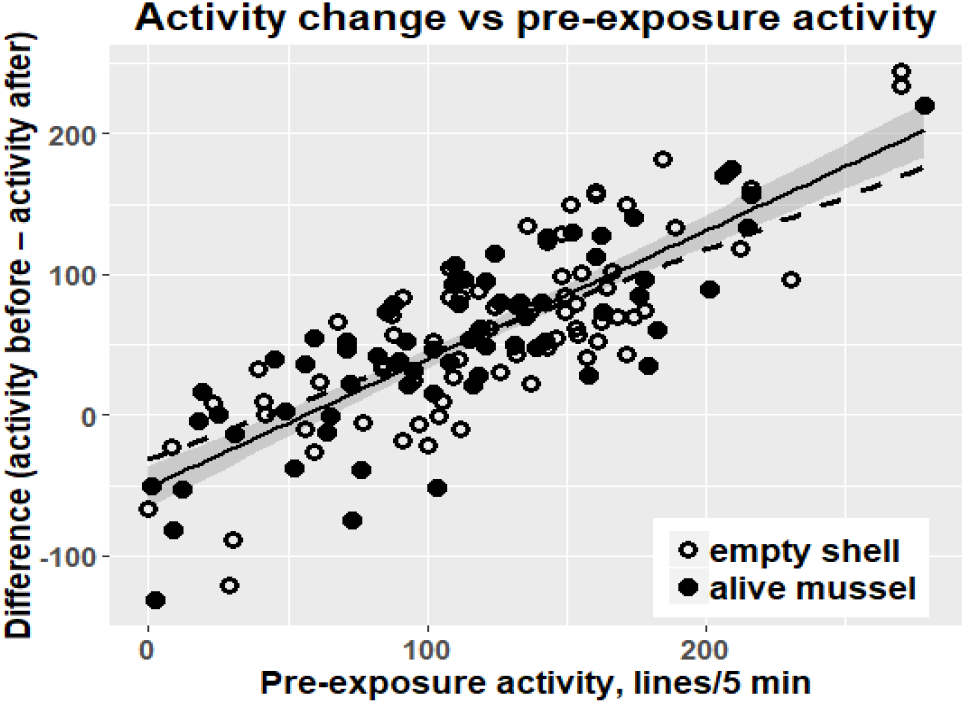
In both treatments fish more active in the pre-exposure period decreased their activity more compared with less active fish (*r* = 0.81, *z* = 3.86, *p* = 0.0001). The solid line is the empirical regression line (*beta* = 0.91), while the dashed line is the regression line fitted using the regression coefficient (*beta =* 0.75) calculated from the null-hypothetical correlation coefficient value.

## Discussion

Though temperature, predation on parasite larvae and host behavior are often reported as important regulators of parasite transmission (Lõhmus & Björklund, 2015; Barber et al., 2016, Barber et al., 2017, Burge et al., 2016, Welsh et al., 2014), their joint effect on transmission success has, to our knowledge, never been tested experimentally.

We found that temperature, presence of filter-feeders (freshwater mussel *A. anatina*) in the environment and individual differences in boldness (activity in the open-field test) had a marked influence on the parasite’s (common eye fluke *D. pseudospathaceum*) infection success. Infection intensity in fish increased with the temperature raise, while the presence of alive mussel led to lower parasitic load in fish. Importantly, there was no interaction between these two factors, which suggests that the effect of freshwater mussel on the infection transmission is constant at least in the temperature range (15-23ºC) tested in our study. Though the increase in the filtration rate with temperature raise was demonstrated at least for several bivalve species under laboratory conditions, the slope of regression curves in these studied were generally gentle (Riisgård & Seerup, 2003; Kittner & Riisgård, 2005)◻◻. Review by Cranford et al. (2011)◻lJ suggested that in natural conditions temperature is unlikely to be an important predictor of the feeding rate in mussels. Filtration rates of mussels usually decrease under high temperatures close to the upper limit of mussel’s physiological tolerance (Ehrich & Harris, 2015; Burge et al., 2016)◻◻. However, water temperatures in our experiment were typical for natural nearshore habitats of *A. anatina* and did not exceed comfort values for this species (Pusch et al., 2001; Falfushynska et al., 2014). Similarly to our results, reduction of trematode transmission by marine bivalves (oysters) was not significantly influenced by temperature, however, the hampering effect of another group of filter-feeders (barnacles) increased with temperature (Goedknegt et al., 2015).

Though *D. pseudospathaceum* cercariae are known to become more infective with temperature (Lyholt & Buchmann, 1996)◻, the mechanism of this phenomenon is unclear. One of the possible explanations for it is the increase of fish motor and ventilation activity with the temperature raise (Krause & Godin, 1995; Pritchard et al., 2001; Mikheev et al., 2014)◻, which is likely to increase host-parasite encounter probability (Barber et al., 2016)◻. Our results showed that correlation between fish motor activity and the temperature was surprisingly weak, however, enhanced ventilation activity, which we did not measure directly, may be responsible for higher infection success under increased temperatures found in our study. An alternative explanation is an immune system function deterioration with temperature increase (Dittmar et al., 2014). Since our heat wave was short-term, it is unlikely to have a strong influence on fish immunity, however, a performance of the innate immunity providing a defense against *D. pseudospathaceum* infection (Scharsack & Kalbe, 2014)◻ deteriorates under warm conditions almost immediately (Dittmar et al., 2014). Another possible explanation is increased activity or/and metabolism of cercariae in warmer water, which can lead to a decrease in cercariae survival (Pechenik & Fried, 1995; Morley et al., 2001), but at the same time can enhance parasite’s infectivity (Poulin, 2006)◻, presumably due to a short-term increase in parasite’s host searching activity and penetration success during short time period.

Only fish activity before the exposure to parasites positively correlated with the parasitic load in fish, while post-exposure activity did not. Fish activity before the exposure strongly varied among fish and, therefore, the difference in motor activity can be a substantial factor explaining differences in parasitic load among individuals. After the exposure, most of the fish decreased their activity, reducing the risk of further infection (Karvonen et al., 2004b; Stumbo et al., 2012). These results are dissimilar with the study on tadpoles, where no significant relationship between pre-exposure activity and infection intensity was found, while activity after the exposure was negatively correlated with parasitic load (Koprivnikar et al., 2012)◻◻. Previously, a decrease in fish activity was reported as a possible defense against a parasitic threat (Stumbo et al., 2012)◻. On the other hand, decrease in activity can also be a non-specific response to the presence of the alarm substances released from the skin of rainbow trout (Sovová et al., 2014)◻ damaged by penetration of *D. pseudospathaceum* cercariae (Poulin et al., 2005). In general, after the exposure to parasites, more active fish decreased their activity stronger than less active ones, which eventually reduced variability in activity after the exposure to parasites. In non-risky conditions, bolder (more active) fish may benefit from quicker food and shelter search, etc., while the main advantage of shyness is lower vulnerability to new threats in the environment. Therefore, when parasite threat arises, activity reduction to some optimal level may become a more beneficial strategy. This Our observation is in conformity with a previous study, which showed that more bold individuals can compensate risky lifestyles with a quicker and more pronounced behavioral response to the parasitic threat (Klemme & Karvonen, 2016)◻. Therefore, we suggest that the fish personality affects fish vulnerability to parasites immediately after the host encounter with the parasitic threat. Later, all fish decrease their activity to a more or less uniform level. Therefore, under common environmental threats, animal personality traits can become less expressed and behavior more uniform shrinking to some optimal level. In other words, fish manifest a kind of a behavioral oddity decrease in risky environments. Though oddity is a well-known factor increasing individual’s susceptibility to predators (Milinski, 1977, Quattrini et al., 2018, Rodgers et al., 2015), it was mainly considered from the predator’s point of view. However, animals’ ability to self-tune their personality traits in risky environments to avoid other potential threats, e.g. parasites, deserves more attention.

The presence of alive mussels in the environment can influence the relationship between temperature and fish activity. Interestingly, in the absence of alive mussels fish post-exposure activity even decreased with temperature raise, but when mussels were present in the container, the positive relationship between temperature and activity remained. It means that in a more risky environment (without mussels, filtering cercariae) fish may try to compensate for the increased risk of being infected at higher temperatures by changing their behavior in a more radical way. Previously, it was shown that fish, which are less resistant against parasites can invest more in developing parasites avoidance behavior compared with less vulnerable fish (Klemme & Karvonen, 2016).

Our research is likely to reveal only short-term ecological effects of heating within the limits of individual plasticity of studied organisms, while global warming may cause prolonged evolutionary processes, which also should be taken into account.

## Conclusions

In our study, we showed that under temperature increase similar to the predicted for aquatic habitats by the end of this century, fish became more vulnerable to parasitic infection. Though filter-feeders (mussels) can effectively eliminate cercariae from the water, decreasing the parasitic load in fish, this effect remained fairly constant under a relatively broad range of temperatures. Therefore, it is unlikely that filter-feeders can compensate for the increased spread of infectious diseases with climate change as it was previously suggested (Burge et al., 2016).

At the first encounter with the parasitic threat, increase in vulnerability to parasites can be connected with the increased activity of fish caused by the increased temperature, however, these behavioral changes are unlikely to be the only factor predisposing fish to parasites under higher temperatures. After more prolonged exposure to parasites, fish activity substantially decreased and its influence on parasitic loads disappeared. This decrease in motor activity was temperature-dependent and more pronounced in bolder (more active) fish, which led to lower variability in fish activity in the presence of parasites compared with the safe environment. Therefore, when working together, warming and parasite threat both can influence fish behavior, altering motor activity and making personality traits less expressed.

## Acknowledgments

We want to thank the technical staff of the Konnevesi research station (University of Jyväskylä, Finland) for their assistance. The research was supported by the Academy of Finland (JT, mobility grants 311033 and 326047); the Otto Kinne Fellowship (EM, grant 2017); the Ella and Georg Ehrnrooth Foundation (MG, mobility grant 2018), the Russian Foundation for Basic Research (VM, grant 17-04-00247); the Ministry of Education and Science of the Russian Federation (AP, the state assignment theme 0149-2019-0008) and Russian Science Foundation grant to VM and MG (19-14-00015).

Authors declare that they have no conflicts of interest.

## Authors’ contributions

All authors conceived the study. MG and EM conducted the experiments, performed the statistical analysis and wrote the major part of the article. AP, VN and JT discussed the results of the study, wrote minor passages of the text and revised the manuscript. JT supervised the study.

## Data accessibility

All data used in the paper are stored in the figshare repository and can be accessed freely (https://doi.org/10.6084/m9.figshare.8080907).

