## Supplement for "Parasite transmission in aquatic ecosystems under climate change: joint effects of temperature, host behavior and elimination of parasite larvae by predators"

*Methods*

1. ***Mean summer temperatures*** in temperate lakes were calculated using data from ‘laketemps’ package (Sharma et al. 2015). This database contains information concerning summer lake surface temperatures for 291 lakes collected *in situ* and/or by satellites for the period 1985–2009. For our calculations, we used data measured using satellite methods. We considered lakes situated between 40° and 68° latitude as temperate lakes (91 lakes). The mean summer temperatures for each lake over all years was used for calculations.

***Statistical analysis***

2. To present the results of the mixed-effect models graphically, ***partial regression (=added variable) plots*** were drawn. To create them, the dependent variable (log transformed infection intensity) and the variable of interest (temperature) were regressed against continuous variables (mass and mass + pre-infection activity for full and abridged datasets respectively). Then the residuals from the models were regressed against each other. The influence of categorical predictor (presence/absence of the mussel in the container was shown using two types of dots. ‘Filled’ dots represent data points from treatments with mussels, while ‘empty’ dots are data points from the control treatment. In addition, we plotted models’ estimates with 95% confidence intervals to visualize the magnitude and the range of the regression coefficients.

3. ***Measuring the relationship between baseline level of activity and subsequent change in activity after the exposure*** is not a straightforward task, because of two common problems: mathematical coupling and regression to the mean.

Thus, mathematical coupling problem arises, when one of the correlated variables is a part of another variable (e. g. when we are regressing y versus y – x). It is easy to demonstrate (Chiolero et al. 2013) that even when x and y are series of independent random numbers with the same standard deviation, a strong Pearson’s correlation (expected value ≈ 0.71) between x and x – y is likely to be observed. Therefore, the null hypothesis that the correlation between baseline value and subsequent change equals 0, as for ‘normal’ regression, is incorrect. Moreover, when x and y are somewhat correlated the situation becomes even more complicated since in such cases the null hypothesis concerning the correlation between y and y – x can differ both from 0 and 0.71 (Tu et al. 2005).

Regression towards the mean problem occurs when the variable is extreme at its first measurement but is closer to the mean at its second measurement. For instance, an individual fish activity can have an expected value of *mu*, however, at the first measurement, a researcher by the pure chance obtained *mu* + *3*sigma*, which is, of course, possible though highly unlikely. However, at the repeated measurement on the same individual, it is much more likely to get the activity value much closer to the *mu*. Therefore, more active fish at the first measurement can be less active at the second measurement partly simply by chance, which may result in a positive relationship between baseline level of activity and change in the activity. In our study situation is not exactly similar to the described above since expectations of fish activity before and after exposure are unlikely to be the same *mu*, however, the logic stays the same.

There are several approaches to account for these problems, we used the method suggested by Tu et al. (2005) because it does not contradict the graphical representation of the results. For instance, common Oldham’s method, which gives similar results on our data, suggests regressing y + x vs y – x, rather than y vs y – x.

Following Tu et al., 2005, we calculated a correlation coefficient between pre-exposure activity and the difference between pre- and post-exposure activity (r_x,y – x_). Then, a correct null hypothesis (r_h0_) was determined taking into account correlation (r_xy_) between pre- and post-exposure activity of fish using the following formula:

$r_{h0}=\sqrt{\left( \frac{1-r_{xy}}{2} \right)}$

Since Pearson’s correlation coefficients are not normally distributed, they were z transformed Finally, both observed and expected (null hypothesis) correlation coefficients were z-transformed:

$z_{r}=0.5*ln\left( \frac{1+r}{1-r} \right)$

and a difference between them was compared with 0 using the z-test using the following expression:

$z=\frac{z_{r\left( x,x-y \right)}-z_{r\left( h0 \right)}}{\sqrt{1/\left( n-3 \right)}}$

where r_H0_ and r_x,y – x_ are the H0 correlation coefficient and sample correlation coefficient respectively, while the null-hypothesis tested is that r_H0_ – r_x,y – x_ = 0, which is tested by testing z_r(x,y – x)_ – z_r(h0)_ = 0

4. ***Influence of environmental conditions and fish phenotype on the infection intensity (when the temperature was considered a factor variable).*** All the details of the analysis were similar to ones described in the main text with only one difference. The biologically sensible model of interest included the temperature predictor turned to a factor as it was initially planned when conceiving the experimental design of the study. The factor has three levels: control temperature, mild heating, and strong heating. In three tests, the temperature in containers with heating was set close to 19.5°C (mean±SD = 19.6±1.59°C), while in four others it was around 22.5°C (22.6±1.48°C). In control containers, the temperature was about 15-17°C (16.0±0.70°C).

The model was the following: log(infection intensity) ~ fish mass (covariate) + temperature (factor) + alive mussel presence/absence (factor) + temperature*alive mussel presence/absence + pre-exposure activity + experiment identity (random factor). Since we were interested in certain double interaction (temperature*alive mussel presence/absence) we included only this double interaction in our model of interest. Then, we simplified the model using backward selection tool from the ‘lmerTest’ package (Kuznetsova et al. 2017). P-values were calculated using Kenward – Roger’s procedure for the approximation of degrees of freedom implemented in lmerTest package (Kuznetsova et al. 2017). An addition of the interaction of interest (*F_2, `129.0_* = 1.14, *p* = 0.32; *F_2, `166.0_* = 0.89, *p* = 0.41) and the fish mass (*F_1, `131.4_* = 0.96, *p* = 0.33; *F_1, `168.6_* = 2.50, *p* = 0.12) did not appear to explain substantial amount of variance. However, as it was described in the main text of the article, we decided to leave the mass in the final models, since it seems to be biologically relevant. Therefore, our final models were the following: log(infection intensity) ~ fish mass (covariate) + temperature (factor) + alive mussel presence/absence (factor) + experiment identity (random factor) + pre-exposure activity for the abridged datasets, while for full dataset it was log(infection intensity) ~ fish mass (covariate) + temperature (factor) + alive mussel presence/absence (factor) + experiment identity (random factor), i.e. similar but excluding activity, since videos for one of the experiments were accidentally lost. 'multcomp' package (Hothorn et al. 2008) was used to conduct multiple comparisons of means (Tukey contrasts) between three heating treatments.

5. ***Software used in the analyses***

In addition to the R packages mentioned in the main test, in the Supplement, we also used 'multcomp' package (Hothorn et al. 2008) to conduct multiple comparisons of means (Tukey contrasts) between the three heating treatments.

***Results***

| **Table S1**. GLMM on full and abridged datasets summary tables. Log-transformed infection intensity was a response variable. | | | | | | | | | | |
| --- | --- | --- | --- | --- | --- | --- | --- | --- | --- | --- |
|  | Full dataset models | | | | | Abridged dataset models | | | | |
| Fixed effects | df | *t* | *p*-value | Est. | *SE* | df | *t* | *p*-value | Est. | *SE* |
| + strong heating | 170.4 | 8.23 | **<0.0001** | 0.707 | 0.080 | 132.9 | 7.49 | **<0.0001** | 0.661 | 0.088 |
| + mild heating | 170.8 | 2.45 | **0.015** | 0.226 | 0.092 | 132.2 | 0.93 | 0.36 | 0.105 | 0.114 |
| + alive mussel | 169.0 | -5.02 | **<0.0001** | -0.303 | 0.060 | 131.0 | -4.46 | **<0.0001** | -0.308 | 0.069 |
| + activity (before) |  |  |  |  |  |  | 2.21 | **0.029** | 0.0015 | 0.0007 |
| + fish mass | 169.6 | 1.57 | 0.12 | 0.026 | 0.016 | 131.4 | 0.98 | 0.33 | 0.0196 | 0.020 |
| Multiple comparisons of means: Tukey contrasts | | | | | | | | | | |
| Linear hypotheses | *z*-value | | *p*-value | Estimate | *SE* | *z*-value | | *p*-value | Estimate | *SE* |
| Strong vs control | 8.82 | | **<0.001** | 0.707 | 0.080 | 7.51 | | **<0.001** | 0.661 | 0.088 |
| Mild vs control | 2.45 | | **0.036** | 0.226 | 0.092 | 0.93 | | 0.61 | 0.105 | 0.113 |
| Strong vs mild | -3.956 | | **<0.001** | -0.482 | 0.122 | -3.91 | | **<0.001** | -0.555 | 0.142 |

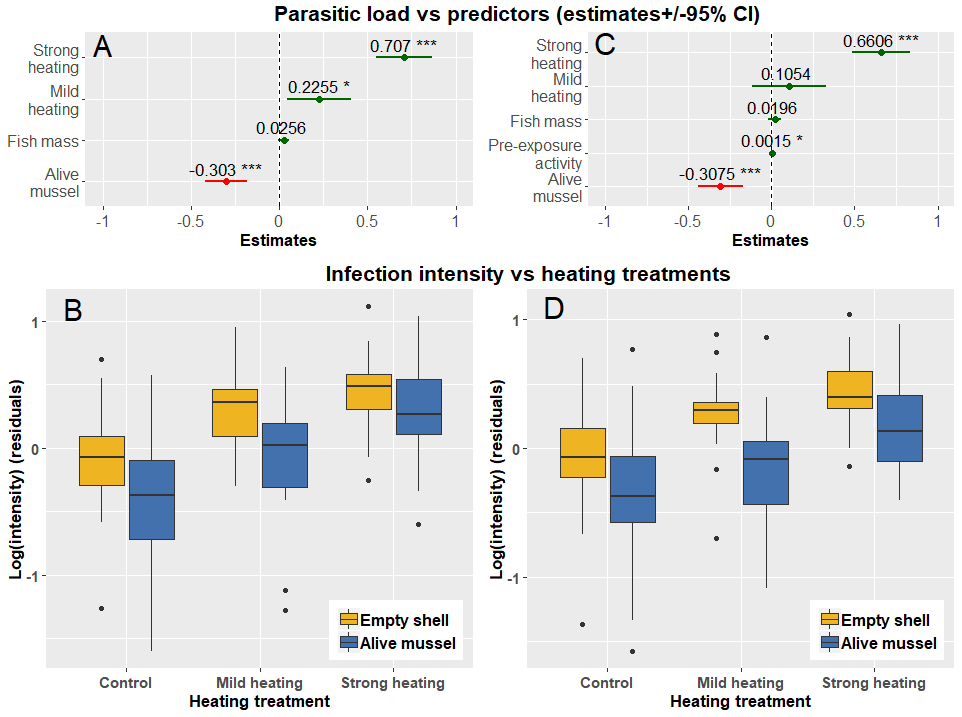
**Fig. S1**. Regression coefficients plots (A, C) and boxplots (B, D) showing the influence of heating treatments on the infection intensity in rainbow trout for the models fitted on the full (A, B) and abridged dataset (C, D). In both cases, the presence of alive mussel in the container substantially (26% and 27% respectively) decreased the infection intensity in fish, while temperature increase led to higher infection intensities. In full dataset both strong and mild heating lead to increased infection intensity in fish (25% and 102% respectively), while in abridged dataset only the influence of strong heating was substantial (94%), most likely because of the smaller sample size in mild treatment in the abridged dataset. There was no significant interaction between the temperature and presence of alive mussel in the environment.
